# Pharmacological validation of an attention bias test for conventional broiler chickens

**DOI:** 10.1101/2024.01.12.575423

**Authors:** Marconi Italo Lourenço da Silva, Alexandra Ulans, Leonie Jacobs

## Abstract

Fear and anxiety are considered concerns for animal welfare as they are associated with negative affective states. This study aimed to pharmacologically validate an attention bias test (ABT) for broiler chickens (*Gallus gallus domesticus*) as a cognitive bias test to determine anxiety. Two-hundred-and-four male Ross 708 broiler chickens were arbitrarily allocated to either the anxiogenic or control treatment at day 25 of age, resulting in 102 birds per treatment. Birds from the anxiogenic group were administered with 2.5 mg/kg of β-CCM (β-carboline-3-carboxylic acid-N-methylamide [FG 7142]) through an intraperitoneal injection at a volume of 0.1 ml/100 g of body weight. Birds from the control group were administered with 9 mg/kg of a saline solution. During ABT, birds were tested in groups of three (n = 34 groups of three birds/treatment) with commercial feed and mealworms as positive stimuli and a conspecific alarm call as a negative stimulus. Control birds were 45 s faster to begin feeding than anxiogenic birds. Birds from the control group vocalized 40 s later and stepped 57 s later than birds from the anxiogenic group. The occurrence of vigilance behaviors did not differ between treatments. This study was successful in pharmacologically validating an attention bias test for fast-growing broiler chickens, testing three birds simultaneously. Our findings showed that latencies to begin feeding, first vocalization, and first step were valid measures to quantify anxiety.

## Introduction

Fear and anxiety manifest as psychological, physiological, and behavioral responses triggered in both animals and humans when there is a perceived or actual threat to their well-being or survival [1,2]. These states are marked by heightened arousal, anticipation, activation of autonomic and neuroendocrine systems, and the display of specific behavior patterns [1]. Fear is a short-term emotional response that drives the instinct to flee or freeze when faced with an imminent, existing threat to survival [3]. Tonic immobility (freezing) elicited by fear is a final defensive strategy observed in prey species, characterized by an anti-predator freezing response or feigning death when captured [4]. Broiler chickens show prolonged freezing in when subjected to rough handling, manual catching compared to mechanical catching, and when exposed to heat stress or shock before testing, in comparison to control conditions [5–8]. In contrast, anxiety is a long-term emotional response that encourages heightened vigilance (i.e., alertness) in reaction to perceived potential threats. This response is intensified by adverse life experiences both before and after hatch [9–11]. These mechanisms have developed as adaptive systems that enhance survival in life-threatening situations by temporarily engaging the sympathetic nervous system and the hypothalamic-pituitary-adrenal axis. This involves suppressing the parasympathetic activity that promotes growth [1]. While anxiety and fear are likely separate emotional states, there may still be some overlap in the underlying brain and behavioral mechanisms [1,3]. It is plausible that anxiety could be considered a more intricate manifestation of fear, offering individuals an enhanced ability to adapt and plan for the future [12].

Animal welfare concerns arise from fear and anxiety as they are negative emotions or associated with negative emotional states [1,13,14]. When these states are consistently activated, they reflect an animal’s struggle to cope with its environment. While fear responses are evolutionarily advantageous for self-preservation, an exaggerated or inappropriate fear reaction can lead to stress, injury, pain, or the development of abnormal behavior, ultimately constituting a negative welfare state [15–18]. Elevated levels of fear and anxiety impair the chicken’ capacity to manage environmental changes, such as handling, restraint, transport, and loud noises [11]. These heightened emotional states have also been associated with a deterioration in feed conversion ratio [5,19].

The attention bias test (ABT) is a cognitive bias test that serves as a tool to determine affective states, particularly anxiety [20]. Within an ABT, an animal’s vigilance or attention toward a perceived threat is assessed [20]. This vigilance can vary and is influenced by affect, wherein heightened levels of anxiety lead to increased vigilance towards a threatening stimulus [14,20]. In the presence of a potential threat, individuals experiencing anxiety tend to exhibit heightened or biased attention directed towards the threat, manifesting as prolonged vigilance and a diminished willingness to feed [2,13,14,21–24]. ABT was pharmacologically validated for sheep [2,24], beef cattle [25], macaques [21], and laying hens [26]. Animals administered an anxiety-inducing drug exhibited heightened attention towards a threatening stimulus and demonstrated increased vigilance compared to their control counterparts [2,21,24–26]. Laying hens that received an anxiogenic drug and were exposed to a conspecific alarm call showed a longer latency to feed, shorter latency to vocalize, and increased locomotion compared to hens injected with saline [26].

Although laying hens are the same species of conventional broiler chickens, these animals are different in terms of body weight, locomotion activity, and emotional reactivity due to divergent artificial genetic selection. Hens are lighter, more active, and show a stronger fear response than broilers [6]. Broilers are prone to impaired gait, difficulty walking, and lameness as they reach slaughter weight and age [27–29]. These physical limitations and the social isolation during an ABT (testing arena) may elicit fear and modify broilers’ responses during the test [14] compared to laying hens.

Chickens have a strong drive for social reinstatement, and in their natural environments, they form highly social groups [30–32]. When pairs of chicks were introduced to a novel open field test, they displayed fewer fear-related behaviors compared to individual chicks undergoing the same test [33]. Slow-growing broiler chickens showed a greater learning success rate in a novel judgment bias test when tested in pairs compared to an individual approach in other studies [34]. Conventional broilers seemed to perform better during an ABT when tested in a group of three compared to broilers tested alone, although no direct comparison was made, and the test approach was not pharmacologically validated [14].

Typically, a pharmacological validation of attention bias towards a threat involves the use of the anxiogenic drug *meta*-Chlorophenylpiperazine (*m*-CPP) [2,24–26,35]. This drug was successfully used to validate ABT in sheep [2,24,35], beef cattle [25], and laying hens [26]. The appropriate dosage of *m*-CPP for broilers is unknown. Another substance used in anxiogenic models to induce anxiety in humans and other animals is the β-carboline-3-carboxylic acid-N-methylamide (β-CCM), also referred to as FG 7142 [36]. A previous study determined that 2.5 mg/kg of body weight was an appropriate dosage of β-CCM for chickens at 25 days old to induce anxiety [37]. The study observed longer durations of tonic immobility, and longer latencies to vocalize and activity in an open field test compared to control chickens. This was interpreted as indicative of heightened levels of fear and anxiety [37]. The current study aims to pharmacologically validate a social-group ABT approach for fast-growing broilers. We hypothesized that broilers injected with the anxiogenic (β-CCM) will vocalize and step faster and will begin to feed later during the ABT compared to broilers from the control group (saline solution).

## Materials and methods

The trial was carried out at Virginia Tech’s Turkey Research Center. Virginia Tech’s Institutional Animal Care and Use Committee approved the experimental protocol (number 22-175) and all procedures were performed in accordance with relevant guidelines and regulations.

### Birds, facilities, and management

Two-hundred-and-four male Ross 708 broiler chickens that were vaccinated for Marek’s disease were obtained at day 21 of age. Birds weighed on average 860±35 g and were sourced from a nutrition experiment performed at the same research center. These birds were part of a larger negative control group and received *ad libitum* access to a standard broiler chicken starter feed as part of their treatment. Throughout that experiment, birds were raised in a fully-automated climate-controlled poultry house. Birds were allocated in pens (1.88 m^2^) with 25 birds each. Pens contained pine shavings, one hanging galvanized tube feeder (SKU# CO30131, Hog Slat, Newton Grove, NC, USA), and one water line with three nipple drinkers (Valco Industries, Inc., New Holland, PA, USA). Calculated stocking density at 21 days was 11.4 kg/m^2^. House temperature was adjusted starting at 34°C on day 1 and reduced approximately 1.5°C every 2-3 days. The birds were maintained on an artificial lighting program of 24L:0D in the first 4 days due to heat lamps and 20L:4D from day 5 to 21 of age.

At 21 days of age, birds were moved to the current study in an almost identical poultry house within the research facility. Here, birds were arbitrarily allocated to 3 pens with 68 chickens each. All pens (8.75 m²) contained pine shavings at approximately 6 cm depth, two hanging galvanized tube feeders, and two water lines, each with three nipple drinkers. Calculated stocking density was 10.0 kg/m^2^ on day 28. Birds had *ad libitum* access to water and pelleted broiler chicken grower feed from day 21 until day 28 (3100 kcal/kg ME and 21.5% CP), and were kept at 18L:6D from day 21 until the end of the trial (day 28), with a light intensity of approximately 15 lux during light hours.

### Experimental treatments and drug injection

Birds were arbitrarily allocated to either the anxiogenic or control treatment at day 25 of age, resulting in 102 birds per treatment. Birds from all three pens were equally distributed across the two treatments. Birds from the anxiogenic group received 2.5 mg/kg of β-CCM (β-carboline-3-carboxylic acid-N-methylamide [FG 7142], Sigma-Aldrich, Merck KGaA, Darmstadt, Germany) dissolved in DSMO (Dimethyl Sulfoxide suitable for HPLC, ≥99.7%, Sigma-Aldrich, Merck KGaA, Darmstadt, Germany) based on Moriarty [37]. Birds from the control group received 9 mg/kg of a saline solution with phenol (0.9% sodium chloride; 0.4% phenol (preservative); *Quantum satis* water for injection; ALK-Abello Pharmaceuticals Inc., Mississauga, Ontario, Canada). Both the anxiogenic and the saline solutions were administered through an intraperitoneal injection at a volume of 0.1 ml/100 g of body weight. Before injection, birds were marked with livestock marker (All-Weather Paintstik, LA_CO Industries, Inc., Elk Grove Village, IL, USA) and weighed to calculate anxiogenic and saline solution dosages. After injection, a 10-minute wait period was applied for both groups so that the anxiogenic solution could take effect.

### Attention bias test

An attention bias test (ABT) was performed by five observers using a modified method as described in [14] over three days when birds were 25, 26, and 27 days of age. The inter-observer agreement was tested for latencies to begin and resume feeding in 21 broilers not included in the current study and was excellent among the five observers (Cronbach’s α of 0.999 for begin feeding and 0.997 for resume feeding).

#### Testing arena

Two square testing arenas constructed with plastic paneling (approximately 119.4 cm L ^x^ 119.4 cm W ^x^ 76.2 cm H) contained pine shavings as bedding and a trough feeder with commercial feed and mealworms. In each arena, two cameras (Teledyne Flir LCC, OR, USA, and EOS Rebel T7 DSLR Camera, Canon, Tokyo, Japan) recorded behavioral responses. The first was placed overhead with a top-down view for live recording and the second provided a side view with recording used for later observations. Both cameras were placed at approximately 1.5 m height. The overhead cameras were connected to an external Lorex NVR (Zhejiang Dahua Technology Co., Ltd., Hangzhou, Zhejiang, China) and two screens in a separate room to allow for undisturbed live behavioral coding. A portable Bluetooth speaker (Fugoo Sport 2.0, Van Nuys, Irvine, CA, USA) was placed near each arena to play a conspecific alarm call at approximately 53 dB. The alarm call signaled a ground predator as in [13,14]. The arenas were located in two separate rooms near the broilers’ home pen room, but far enough to prevent the conspecific alarm call to be heard in the home pens or in the other arena.

#### Testing

Ten minutes after the injection, birds were tested in familiar groups of three (n = 34 groups of three birds/treatment). Each group of three broilers was recorded by one observer. The researcher that injected the anxiogenic or saline solution and their assistant moved birds to and from the testing arena. The observers were blinded to the treatment. After placement, a conspecific ground alarm call was played for 8 s, which is known to elicit a vigilance response in chickens [38]. The alarm call was replayed for 8 s depending on the birds’ behavioral response after the first time the call was played (Table 1). The maximum test duration was 600 s if the alarm call was replayed, or 480 s if it was not.

**Table 1.**
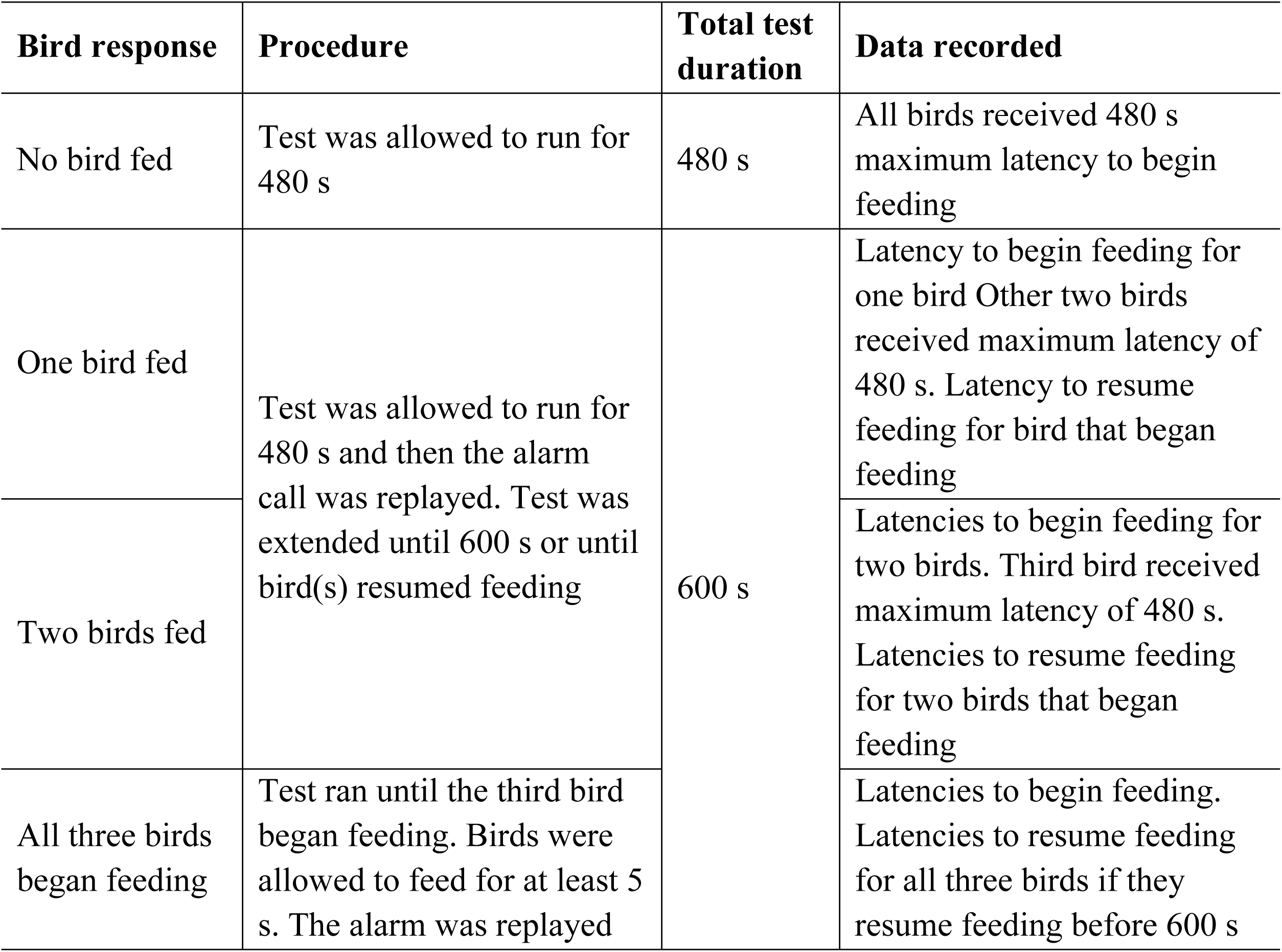
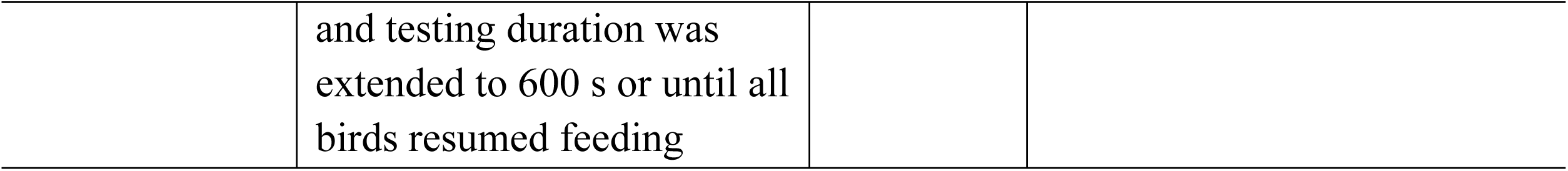
Methods used during attention bias testing after the first alarm call was played. Birds were tested in groups of three. Test duration and data recorded were modified according to the birds’ behavioral responses after the first alarm call was played.

Live-recorded responses included the latency to begin feeding (s) and latency to resume feeding (s) if the alarm call was replayed. Responses collected after the test from video recordings were latency to first vocalization (s, recorded at group level), latency to first step (s), and occurrence (yes/no) of vigilance behaviors (freeze, neck stretches, looking around, and erect posture) in the 30 s following the first alarm call. Each of the four vigilance behaviors was scored as a 1 (occurred) or 0 (did not occur) [14,26].

### Statistical analysis

Data were analyzed in SAS Studio 3.8 (SAS Institute Inc., Cary, NC, USA). Normality of data residuals was assessed by the Shapiro-Wilk’s test. None of the residuals were normally distributed. Generalized linear mixed models (GLIMMIX) were applied for latency to begin feeding, latency to first vocalization, and latency to first step using lognormal distribution, followed by F-tests with treatment as a fixed effect and arena, group, and pen as random effects. No statistical analysis was performed for latency to resume feeding because no birds from the anxiogenic group resumed feeding after the second alarm call was played. Vigilance behaviors (occurrence) were assessed by GLIMMIX using binary distribution, followed by F-tests with treatment as a fixed effect and arena, group, and pen as random effects.

## Results

Latency to begin feeding (mean±SEM) was affected by treatment, with birds from the control group feeding faster than birds from the anxiogenic group (control = 442.6±9.4 s; anxiogenic = 488.0±3.0 s; F_1,138_ = 13.5, *P* < 0.001, Fig 1).

**Fig 1.**
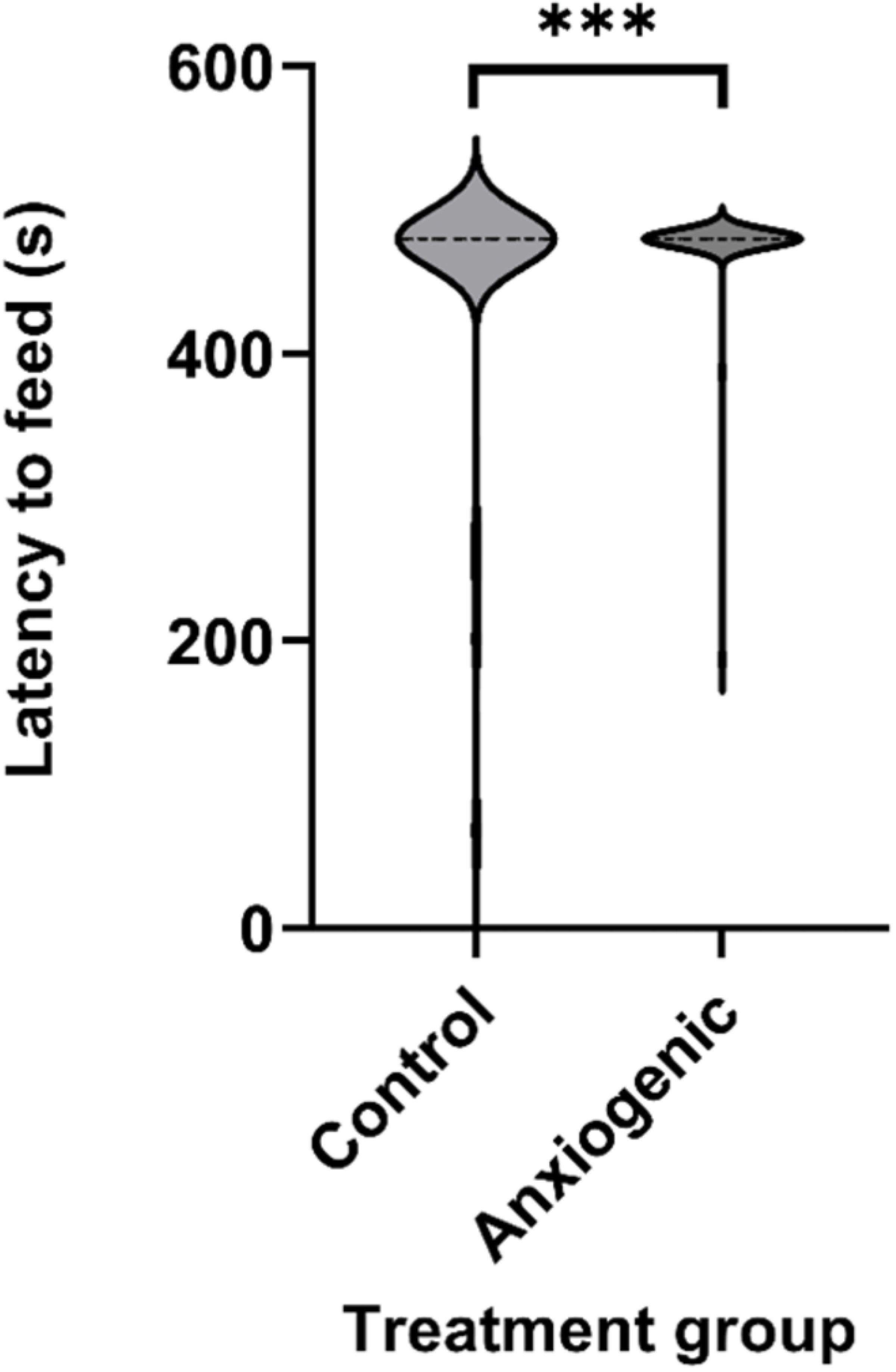
Violin plot of least square mean estimates (s ± SEM) for latency to begin feeding (n = 68 groups of 3 birds) for broiler chickens from control (saline) and anxiogenic treatments at 25, 26, and 27 days of age. *** indicates a difference at *P* ≤ 0.001.

Only two out of ten birds from the saline group that began feeding resumed feeding after the second alarm call (51 and 71 s latencies). No birds from the anxiogenic group that began feeding resumed feeding after the second alarm call.

Latency to first vocalization and latency to first step were affected by treatment. Birds from the control group vocalized later (control = 80.9±10.9 s; anxiogenic = 40.7±5.8 s; F_1,45_ = 13.0, *P* < 0.001, Fig 2) and stepped later (control = 137.6±19.1 s; anxiogenic = 80.8±12.9 s; F_1,29_ = 4.2, *P* = 0.050, Fig 3) than birds from the anxiogenic group. The occurrence of vigilance behaviors did not differ between treatments (*P* ≥ 0.145, Table 2).

**Fig 2.**
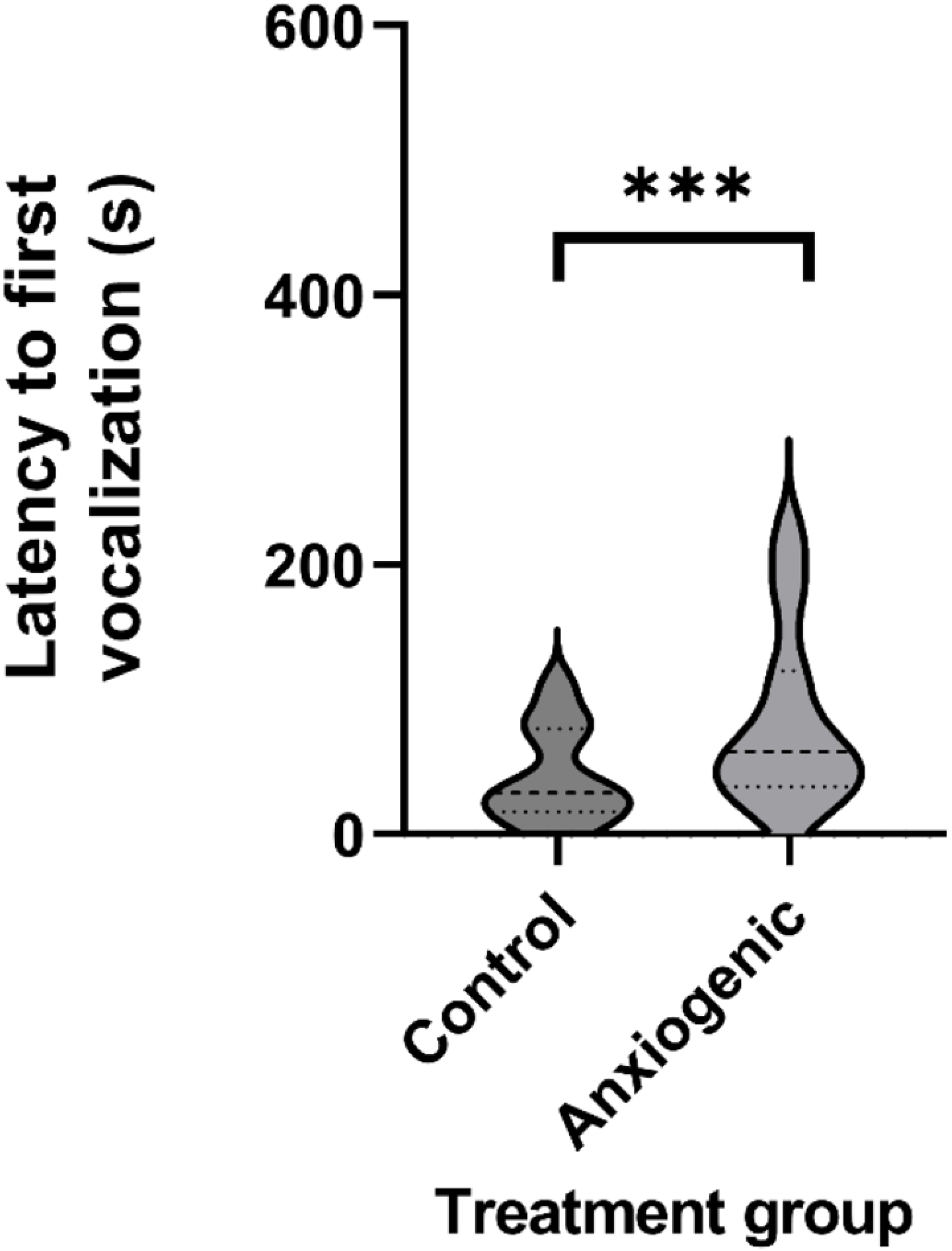
Violin plot of least squares mean estimates (s ± SEM) for latency to first vocalization (n = 23) for broiler chickens from control (saline) and anxiogenic treatments at 25, 26, and 27 days of age. *** indicates a difference at *P* ≤ 0.001.

**Fig 3.**
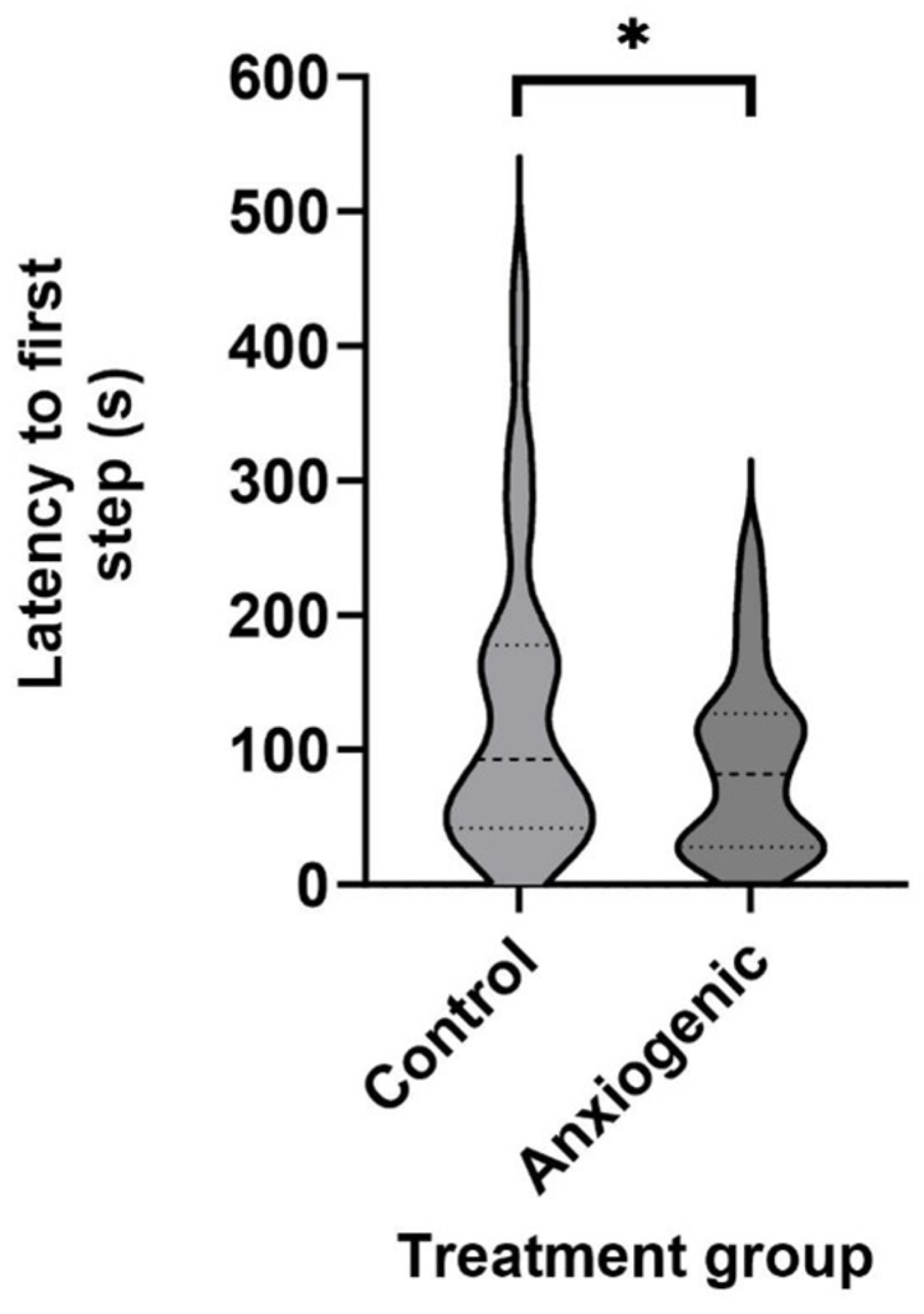
Violin plot of least squares mean estimates (s ± SEM) for latency to first step (n = 33) for broiler chickens from control (saline) and anxiogenic treatments at 25, 26, and 27 days of age. * indicates a difference at *P* ≤ 0.05.

**Table 2.** Proportion (%) of total birds performing vigilance behaviors (n = 68 groups of 3 birds) from either control (saline) or anxiogenic treatments at 25, 26, and 27 days of age.

| Vigilance behaviors | Treatment |  | Test statistic and *P*-value |
| --- | --- | --- | --- |
|  | Control | Anxiogenic |  |
| Erect posture (% of birds) | 44.5 | 46.5 | $F_{1,45} = 0.07$, $P = 0.787$ |
| Neck stretching (% of birds) | 5.9 | 11.1 | $F_{1,45} = 1.67, P = 0.199$ |
| Looking around (% of birds) | 5.0 | 8.1 | $F_{1,45} = 0.18, P = 0.672$ |
| Freezing (% of birds) | 96.0 | 99.0 | $F_{1,45} = 1.65, P = 0.201$ |

## Discussion

This study pharmacologically validated an attention bias test to assess affective state, more specifically anxiety, in broiler chickens. The ABT demonstrated a behavioral distinction between non-anxious, calm chickens (control group) and anxious chickens (anxiogenic group). In order to induce anxiety in chickens, we administrated β-CCM via intraperitoneal injection prior to testing, followed by a 10-min wait period [37]. Previous studies have shown that β-CCM, also referred to as FG 7142, is a validated substance to induce anxiety in chickens and other species [36,37].

Non-anxious, calm chickens in the control group fed faster than chickens from the anxiogenic treatment group, likely because of a stronger septohippocampal-behavioral-inhibition (SBI) system response in the anxiogenic group. When presented with an alarm call threat, individuals that received an anxiogenic biased their attention toward the negative stimulus compared to the positive stimuli (mealworms and food), which aligns with our hypothesis. This attention bias in anxious birds could be associated with activation of the SBI system, which is one of the three primary emotional systems alongside the behavioral approach system and the fight/flight response [39,40]. Animals will compare between actual and expected stimuli in any given situation. The SBI system is activated if there is a discrepancy between those, if the stimulus is predicted to be aversive, for instance if there is a threat of punishment or failure, or if there is uncertainty about the valence of the stimulus. This system activation will increase attention to novel, potentially aversive stimuli and will inhibit the expression of other normal behavior [1]. In the current study, we observed this increased attention towards the presented novel stimuli (alarm call and arena) and the inhibition of normal feeding behavior especially in the anxiogenic group, and less pronounced in the control group. This finding supports that the SBI system was activated in most if not all birds, but more strongly in the anxiogenic treatment group.

Our findings align with pharmacological validation studies of ABT in laying hens [26], sheep [2,24,35], and beef cattle [25]. In this study, chickens injected with β-CCM (the anxiogenic treatment) showed longer latencies to vocalize and step following the first alarm call than chickens from the control treatment. These responses are similar to chickens injected with β-CCM exhibiting a longer latency to vocalize and exhibiting restless behavior in an open field test compared to chickens injected with water [37]. Contrarily, laying hens injected with a anxiogenic drug (*m*-CPP) did not differ from the control group for latencies to first vocalization and first step following the first alarm call in an ABT [26]. The anxious hens in that study vocalized sooner than control hens following the second alarm call, but latencies to first step did not differ [26]. Differences between that and the current may be because they tested hens individually rather than in groups of three. The hens were socially isolated which impacts vocalizations [41,42]. Social isolation may cause them to vocalize sooner than when tested in a group as chickens have a strong motivation for social reinstatement and will vocalize in an attempt to find flock mates [30–32,41,42]. In line, fast-growing broilers performed better (less anxious) in an ABT when tested with two conspecifics compared to being tested alone, although not directly compared [14]. Our results confirm that chickens benefit from social support in testing environments that require attention. Our ABT approach may have improved broilers’ performance during ABT by attenuating the effects of social isolation.

Administering β-CCM in 25-day-old chickens induced anxiety. β-CCM made them restless (stepped and vocalized sooner than chickens from the control group) which could be interpreted as distress in the arena. These behavioral responses are similar to behavioral and physiological responses in other species that were administered with β-CCM, such as speaking (vocalizations), increased plasma cortisol concentrations, and decreased cerebral blood flow in humans [43,44], monkeys [45], mice [46,47], rats [48–50], cats [51], and planaria [52]. Broiler chickens administered with β-CCM remained immobile and displayed longer latencies to vocalize, fewer vocalizations, longer latencies to become active, and lower activity levels than the control group during a tonic immobility test [37] consistent with the hypothesis that β-CCM has anxiogenic properties that induce fear- and anxiety-related behavior [37]. These behavioral changes could be caused by β-CCM acting as a partial inverse agonist at the benzodiazepine allosteric site of the α-GABA_A_ receptors [50]. β-CCM activates neural networks underlying the anxious response [53] and interacts with serotonergic, dopaminergic, cholinergic, and noradrenergic modulatory systems [50]. Our findings indicate that β-CCM did induce anxiety in broilers, as in previous studies.

Broilers and hens show distinctive behavioral differences during the ABT. Broilers in the current study showed different behavioral responses compared to hens and broilers previously. Our broilers’ latency to first vocalization (81 s for control and 41 s for anxiogenic group) was shorter than in laying hens, which vocalized after 110 s (control) and 78 s (anxiogenic group). In contrast, latency to first step was longer (138 s for control and 81 s for anxiogenic group) than previously reported for laying hens (42 s for the control and 52 s for the anxiogenic group) [26]. The differences in ABT responses between broilers and laying hens may stem from diverging genetic strains or ages (developmental stages) at time of testing, as fear is also age-dependent [54–56]. It is widely acknowledged that different strains of domestic fowl exhibit diverse temperaments, particularly in terms of fearfulness or tendencies to flee [57–59]. These may suggest a similar effect of genetics and age on anxiety, although the difference between genetic strains or ages have not been compared directly.

The previous experiences of our sampled birds in their first weeks of life may have shaped their responses in the study. We used broilers from a control group in a nutrition study in order to optimize resources and reduce the use of animals in experiments. It is possible that this prior experience impacted behavioral responses in the current study, as the birds were tested four days after being moved to another facility. All birds underwent the same treatment in the prior study. Individual differences between birds, for instance in cognitive styles [60,61], could have resulted in variation in how these birds experienced that control treatment, yet we anticipated that this would not have a measurable impact on the overall response to their treatment (either anxiogenic drug or control injection). This is supported by the significant treatment effect on test outcomes. From a stress parameter perspective, a 4-day rest period before testing is an appropriate acclimatation period for broiler chickens. Twelve hours’ rest period was enough for broilers to have body temperature, erythrocyte, leukocyte, heterophile, and lymphocyte counts return to baseline after transportation [62]. However, they did not evaluate parameters associated with affective states, thus it is unknown whether affective states returned to baseline. Yet, our study allowed for a longer period of adjustment, and birds were not purposefully stressed prior to transitioning to the current study.

The handling, injection, and wait period at time of testing likely also affected behavioral responses during the ABT, accounting for differences between the current and previous research. Latencies to first vocalization and first step were longer in broilers in the current trial than in broilers from the same genetic strain in our earlier work (vocalize: 15-20 s; step: 27-114 s) [14]. This difference is likely because those broilers did not undergo an injection procedure or prolonged wait period prior to testing [14]. This could explain the behavioral responses indicative of increased anxiety in the current study compared to the previous work. Furthermore, half of the broilers in the previous study were housed in highly complex conditions, which reduced their anxiety during the ABT [14]. Contrary to our predictions, the treatment did not impact the occurrence of vigilance behaviors. In both treatments, freezing behavior was observed in over 95% of the tested chickens within 30 s following the first alarm call. This freezing behavior is likely reflecting an innate fear response [63] caused by the alarm call. Birds diverting their initial attention towards the alarm call, regardless of treatment, suggests that they perceived the alarm call as a potential threat [26]. The lack of difference in initial responses between treatments suggests that all birds’ initial response to the alarm call was fear, followed by attention towards the potential threat. This attention was prolonged in the anxiogenic treatment group compared to the control group, as reflected in the longer latency to begin feeding than birds in the control group. Responses in our study are consistent with results in sheep that showed similar vigilance behaviors when comparing the control and anxiogenic treatment groups [24].

The handling, injection, and wait period at time of testing may also explain why vigilance between treatments did not differ. Fear induced by the injection procedure and the prolonged wait period may have caused vigilance to be similar in both treatments. Distressing situations such as handling, restraint, transport, and injections cause fear in production and laboratory animals [6,11,64,65]. This fear response in both treatments may be linked to the anti-predatory response in chickens. Chickens are precocial [66,67], exhibit anti-predatory and avoidance behaviors already early in life [68,69]. They do not need previous experience to flee or freeze in response to a predator threat [63]. The stress response associated with veterinary procedures, such as injections, can vary among individual animals, with one experiencing substantial distress while another may be less impacted [64]. The intensity of a physiological or behavioral response during handling is influenced by prior experiences [11,70–72]. The chickens could have experienced the injection procedure as aversive handling since even using a gentle approach, they were restrained and inverted while receiving an intraperitoneal injection. It might have elicited anti-predatory and fear response [73]. Therefore, handling prior to the ABT could have increased vigilance in all tested birds regardless of treatment.

Vigilance behavior in the first 30 s of testing may not be a valuable measure of anxiety during the ABT. Vigilance behaviors during ABT were inconsistent across studies. In beef cattle and sheep, anxiety-induced animals showed more vigilance behaviors compared to the control group [2,25,35]. Hens housed in cages or enriched pens did not show differences in vigilance behaviors among treatments [13]. Hens injected with a saline solution did not differ in vigilance behaviors compared to hens injected with an anxiogenic solution [26]. More broilers housed in low-density pens were vigilant (looking around) than those housed in high-density pens, while other vigilance behaviors did not differ [14]. Because these inconsistencies, we propose to use this measure to indicate that the alarm call induced anxiety, rather than as a measure to compare between treatments. Alternatively, lengthening the time of observation of vigilance behaviors or quantifying durations rather than occurrence may results in more consistent contrasts between treatments.

To conclude, this study was successful in pharmacologically validating an attention bias test for fast-growing broiler chickens, testing three birds simultaneously. Our findings showed that latencies to begin feeding, first vocalization, and first step were valid measures to quantify anxiety, as chickens from the anxiogenic treatment group exhibited longer latencies to begin feeding, and shorter latencies to first vocalization and first step compared to the control group. Vigilance behaviors in the first 30 s of the test period were not valid indicators of anxiety, but rather indicative of a state of fear or anxiety directly after the initial alarm call was played. The attention bias test can be used to infer an affective state in broiler chickens, more specifically anxiety. This test could be used to inform industry decisions related to management and housing conditions, and their impact on a key aspect of broiler chicken welfare. The test shows promise to be applied in commercial conditions, for instance as part of an animal welfare audit.

## Supporting information

**S1 Fig 1. Violin plot of least square mean estimates (s ± SEM) for latency to begin feeding (n = 68 groups of 3 birds) for broiler chickens from control (saline) and anxiogenic treatments at 25, 26, and 27 days of age. *** indicates a difference at *P* ≤ 0.001.**

**S2 Fig 2. Violin plot of least squares mean estimates (s ± SEM) for latency to first vocalization (n = 23) for broiler chickens from control (saline) and anxiogenic treatments at 25, 26, and 27 days of age. *** indicates a difference at *P* ≤ 0.001.**

**S3 Fig 3. Violin plot of least squares mean estimates (s ± SEM) for latency to first step (n = 33) for broiler chickens from control (saline) and anxiogenic treatments at 25, 26, and 27 days of age. * indicates a difference at *P* ≤ 0.05.**

**S1 Table 1. Methods used during attention bias testing after the first alarm call was played. Birds were tested in groups of three. Test duration and data recorded were modified according to the birds’ behavioral responses after the first alarm call was played.**

**S1 Table 2. Proportion (%) of total birds performing vigilance behaviors (n = 68 groups of 3 birds) from either control (saline) or anxiogenic treatments at 25, 26, and 27 days of age.**

